# Cradle to production gate life cycle assessment of cultured meat growth media: A comparison of Essential 8™ and Beefy-9

**DOI:** 10.1101/2023.04.21.537772

**Authors:** Derrick Risner, Patrick Negulescu, Yoonbin Kim, Cuong Nguyen, Justin B. Siegel, Edward S. Spang

## Abstract

There is an increasing interest in use of biotechnology as a means of sustainable food manufacturing; however, biotechnology processing is considerably resource and energy intensive. Recent interest in animal cell-based meat (ACBM) has prompted scientific and engineering questions about the economic and environmental viability of these proposed ACBM products. This study provides an environmental assessment of two proposed growth mediums (Essential 8^TM^ and Beefy-9) for ACBM production. The study found that the addition of antibiotics/antimycotics (10,000 μg/mL) to the growth media increased the environmental metrics, such as the cumulative energy demand and global warming potential, by two orders of magnitude. To account for additional processing for animal cell culture, a scenario analysis was conducted to assess the potential environmental impacts of growth medium production with varying level of refinement required for the input components. The study indicates that the heavy refinement of the growth medium components is likely to undermine the potential sustainability of future ACBM products.

## 1. Introduction

Biotechnology has historically been utilized to preserve and enhance food properties and is currently responsible for many advancements in the pharmaceutical and bioindustrial sectors. The modern bio-based economy includes not just food and animal feed, but also bio-based chemicals, materials, health products, and bio-based fuel (Lange et al., 2021). Bio-based economies have been prescribed as sustainable and as a potential means to achieve many of the United Nations’ Sustainable Development Goals (SDG) (Lange et al., 2021; United Nations, 2015). More specifically, in response to SDG 2 (End hunger, achieve food security and improved nutrition, and promote sustainable agriculture), bioprocessing technology utilizing bioreactors has been proposed for food/protein/meat production (Moritz et al., 2015; Post, 2012; E. A. Specht et al., 2018; L. Specht, 2019). Additionally, there has been sizable financial investment (>1 billion USD) in companies which aim to utilize bioreactor-based technology to produce animal cell-based meat or “cultured meat” (Risner et al., 2020; Turi, 2021).

In contrast, critics have raised concerns that a strong focus on developing a bio-based economy may actually hinder the achievement of some of the ecological SDGs (Fritsche & Iriarte, 2014; Heimann, 2019). For example, while pharmaceutical technology has produced many positive impacts for human health, it has been reported that the global pharmaceutical industry has a greenhouse gas (GHG) emissions intensity 55% higher than the automotive industry (Belkhir & Elmeligi, 2019; Buxbaum et al., 2020). It has also been reported that the production of active pharmaceutical ingredients has a cumulative energy demand twenty-times greater than bulk chemical production (Wernet et al., 2010). These results indicate that additional efforts to critically examine the proposed biotechnological solutions for the SDGs are required to inform environmentally attuned decision-making and investment in this space.

Previous efforts to quantify the environmental impact of cultured meat have been based on forward-looking projections and do not entirely account for all the inputs and processes required for animal cell culture (Mattick et al., 2015; Tuomisto et al., 2014; Tuomisto & Teixeira de Mattos, 2011). A gap analysis of existing life cycle assessments (LCAs) of cultured meat specifically identified the need for a more robust environmental assessment of the animal cell growth media, i.e. the nutrient-dense broth in which the target cells are cultivated (Carus et al., 2019). Previous work also indicates that the volume of media required for industrial cultured meat production is a limiting economic factor (Risner et al., 2020). This indicates that quantification of the embedded resources within the animal cell growth media is necessary to evaluate the environmental impact of this potential food production technology.

Growth media for animal cell culture can vary in composition but can be broadly categorized as either “complex” or “defined” growth media. Complex media is inherently variable, containing components which are not completely chemically defined such as fetal bovine serum. This can introduce unknown factors which can affect animal cell proliferation and differentiation. The utilization of growth media containing animal-based components would also largely be contradictory to the “spirit” of the cultured meat products, especially in terms of the technology addressing the issue of animal welfare in conventional meat production system (Lanzoni et al., 2022). In contrast, defined media is chemically defined with set concentrations of proteins, amino acids, sugars, vitamins, minerals, salts, among other constituents.

Essential 8^TM^ (E8) is a defined growth medium which has been utilized and promoted as a viable growth medium for stem cells and ACBM production (Chen et al., 2011; Kolkmann et al., 2020; L. Specht, 2019; Verbruggen et al., 2018). The E8 growth medium was originally designed for researchers studying human induced pluripotent stem cells and embryonic stem cells. E8 was formulated as a consistent, defined medium to improve experiment reproducibility, but was not originally designed as a growth medium for industrial cell biomass production (Chen et al., 2011).

E8 is largely composed of Dulbecco’s Modified Eagle Medium/Hams’ F12 (DMEM/F12) basal medium, which is widely used for animal cell culture along with 7 other ingredients, including: 2-phospho-L-ascorbic acid trisodium salt, insulin, transferrin, sodium selenite, fibroblast growth factor-2 (FGF-2), transforming growth factor beta (TGF-β), and additional sodium bicarbonate. DMEM/F12 is also the base for the recently developed animal cell growth medium, Beefy-9 (B9) (Andrew Stout et al., 2021). In addition to DMEM/F12, B9 contains the same components as E8 with the additional components of neuregulin, ultrapure water, antibiotics/antimycotics, and recombinant albumin (Stout et al., 2021).

In sum, understanding the environmental impacts of producing E8/B9 animal cell growth media requires identifying tracking, and consolidating the embedded resources utilized and waste outputs generated in the production of each media ingredient. The analysis assumes that all medium components are produced and purified individually and then mixed in the appropriate proportions during the media preparation. This is no small task considering that both E8 and B9 growth media are composed of more than 50 different input ingredients when DMEM/F12 is broken down into its constituent components. Thus, to understand the potential environmental impact of E8/B9 production, we include all these ingredients (or at least as many as could be included given data availability) in our comparative life cycle assessment (LCA).

## 2. Materials and methods

All of the individual components of the E8/B9 media were identified and categorized into eight broad categories (Figure 1.0) based upon their production method (Chen et al., 2011; L. Specht, 2019). Table 1.0 provides the composition of each growth media by their component ingredients. The LCA was conducted following the ISO 14040 and 14044 standards to estimate the potential environmental impact of each E8/B9 component (International Organization for Standardization, 2006a, 2006b). A combination of peer-reviewed literature, OpenLCA v.1.10 software, existing databases, stoichiometric calculations, and engineering judgment was utilized to understand each E8/B9 component production process and to complete the LCA.

**Figure 1.0.**
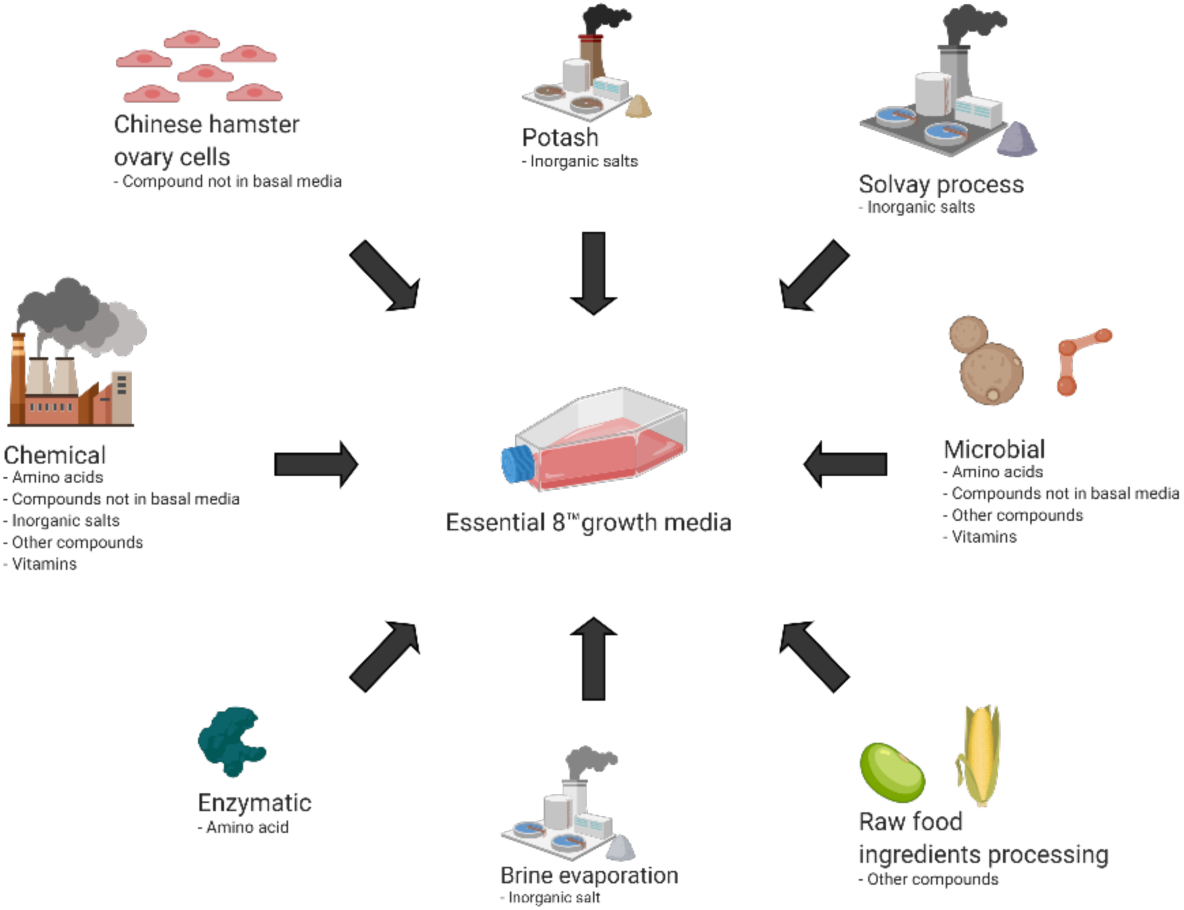
Broad production categories for E8/B9 components

**Table 1.0.** Essential 8 and Beefy-9 growth medium composition

| Component | Concentration in Essential 8™ (µg/mL) | Concentration in Beefy-9 (µg/mL) |
| --- | --- | --- |
| DMEM/F12 basal media | - | - |
| 2-Phospho-L-ascorbic acid trisodium salt | 64 | 200 |
| Insulin (human, recombinant) | 19.4 | 20 |
| Transferrin (human, recombinant) | 10.7 | 20 |
| Sodium selenite | 0.014 | 0.02 |
| Fibroblast growth factor (FGF-2) | 0.1 | 0.04 |
| Neuregulin (NRG1) | - | 0.0001 |
| Transforming growth factor (TGF-β3) | 0.002 | 0.0001 |
| UltraPure Water | - | 58000 |
| Antibiotic/Antimycotic | - | 10000 |
| Recombinant albumin | - | 800 |
| Additional NaHCO<sub>3</sub> | 543 | - |

### 2.1 Goal and Scope

The goal of this LCA was to estimate the environmental impact of current/near-term E8 and B9 growth medium production. It is hoped that the LCA results will provide clear environmental impact information to producers utilizing large volumes of E8/B9 growth mediums or other mediums produced in a similar manner. The analysis was conducted as a cradle-to-production gate analysis. A system boundary was set at the cradle (raw material extraction) to the E8/B9 production facility gate. A limit of 0.1 kg reactant or precursor per kilogram of input was deemed the minimum limit to continue to track a component. For the sake of this study, precursor refers to a material/chemical used to produce an ingredient in E8/B9 growth medium (ex. starch hydrolysate is a precursor to glucose). The functional unit was defined as a liter of growth medium with the reported concentrations of each component (L. Specht, 2019; Stout et al., 2021). The functional unit was chosen to allow for comparison with other defined cell growth mediums.

### 2.2 Life cycle inventory (LCI)

Production process information for each medium component was initially searched for in the ecoinvent (v.3.8) database. If available, then the material and energy input flows were tracked utilizing the ecoinvent (v.3.8) datasets (ecoinvent Association, 2021). If the initial production process information was not available in ecoinvent, then other literature sources and calculations were utilized to estimate material inputs and outputs (see following section and Appendix A-H). Ecoinvent’s global datasets were utilized throughout the life cycle inventory to limit the effect of geographic variation. The ecoinvent database can be examined with five different settings (undefined, allocation (cutoff by classification), allocation at the point of substitution, substitution (consequential, long term), and allocation (cut-off, EN15804)) which unlink or link datasets using several different methodologies. The database search was configured to “undefined” to maximize the LCI analysis transparency (ecoinvent Association, 2021). An undefined system model unlinks unit processes and allows for multiple outputs from each unit process. The flows and processes were then imported and configured in OpenLCA software which tracks inputs/outputs for a product system. The estimated material and energy flows should be considered non-exhaustive as the industrial production processes for some medium components (e.g. 4-(2-hydroxyethyl)-1-piperazineethanesulfonic acid (HEPES) and lipoic acid production) were excluded and other E8/B9 component production processes were only partially represented as a result of gaps in the available data. It should also be noted that the reported E8/B9 component production processes do not represent the production of cell culture-grade materials. Production of more highly purified cell culture-grade materials require additional resources, and this is addressed in the scenario analysis section.

The methods, calculations, limitations, and assumptions for the LCA model are further elaborated in the subsequent sections, including additional detail on each of the eight production categories identified in Figure 1.

#### 2.2.1 Raw Food Ingredients

Corn was assumed to be the source for glucose due to to the fact that it is already widely utilized for biorefining and food/beverage production in the United States (Capehart & Proper, 2021).

Cottonseed oil production was utilized to estimate linoleic acid production due to its alignment with the cottonseed oil fatty acid profile (Yang et al., 2019). Ecoinvent datasets were utilized to estimate the material flow for both glucose and linoleic acid. Appendix A provides details on the calculations and procedures utilized to determine the material flows of glucose and linoleic acid.

#### 2.2.2 Microbial Fermentation Products

Components of E8/B9 which are, or have potential to be, produced via microbial fermentation were identified (Tables A1.0 and A2.0). The total mass of each component was determined from literature (Chen et al., 2011; L. Specht, 2019; Stout et al., 2021). The glucose mass requirement for each component was determined utilizing microbial yields (g product/g glucose) and microbial titers (g/L of media) from literature sources (see appendix B). Microbial yields with greater than 0.01 g product/g glucose were utilized (if available in literature), since the glucose concentration can vary depending on organism growth requirements, fermentation system, and operating parameters (Wu & Maravelias, 2018).

When a microbial yield was unavailable for a growth medium component, microbial titers (g/L) from the literature were utilized to estimate the required mass of glucose. The glucose concentration of the media was assumed to be 10 g/L for calculations which utilized titer to estimate the required glucose mass. A batch system without the capabilities to add supplemental nutrients/glucose was assumed. Given this assumption, a glucose concentration of 10 g/L was deemed acceptable (Millipore Sigma, n.d.).

The inputs/outputs other than glucose for microbially-produced compounds were estimated utilizing data from industrial lysine production as a proxy system. Varying yields between compounds indicated that a correction factor was necessary, i.e. more resources are utilized if more batches are required for the same mass of product. Each correction factor was calculated utilizing the reported lysine yield and the reported compound yields (Marinussen & Kool, 2010). When microbial titer was reported and utilized in the model, an assumed glucose concentration (10 g/L) was used to calculate the correction factor. Table A1 and A2 in appendix B provide correction factors and sources for yields and titers (See calculations A2 and A3 in appendix B).

#### 2.2.3 Enzyme-derived Products

The embedded resources for the enzymatic production of E8/B9 components were estimated utilizing a similar approach as previously described in microbial fermentation products section. L-aspartic acid was the only E8/B9 component identified to be produced enzymatically and the description of the assumed process can be found in Appendix C.

#### 2.2.4 Chemical Products

The ecoinvent database was utilized to estimate embedded energy and material flows for compounds produced via chemical synthesis (ecoinvent Association, 2021). If the ecoinvent datasets were not available, reported production methods for the compounds were analyzed and stoichiometric calculations were conducted to determine the mass of E8/B9 component precursors (reactants). This process was repeated if the E8/B9 precursor was not available in the ecoinvent dataset. Figure 2.0 provides an example of this process with the components encased in the red outline having datasets available in ecoinvent and these datasets are utilized to account for the environmental impact of each component. Substitution was also utilized if the data were unavailable in the ecoinvent dataset for particular E8/B9 components (e.g. ascorbic acid was substituted for ascorbic acid 2-phosphate).

**Figure 2.0.**
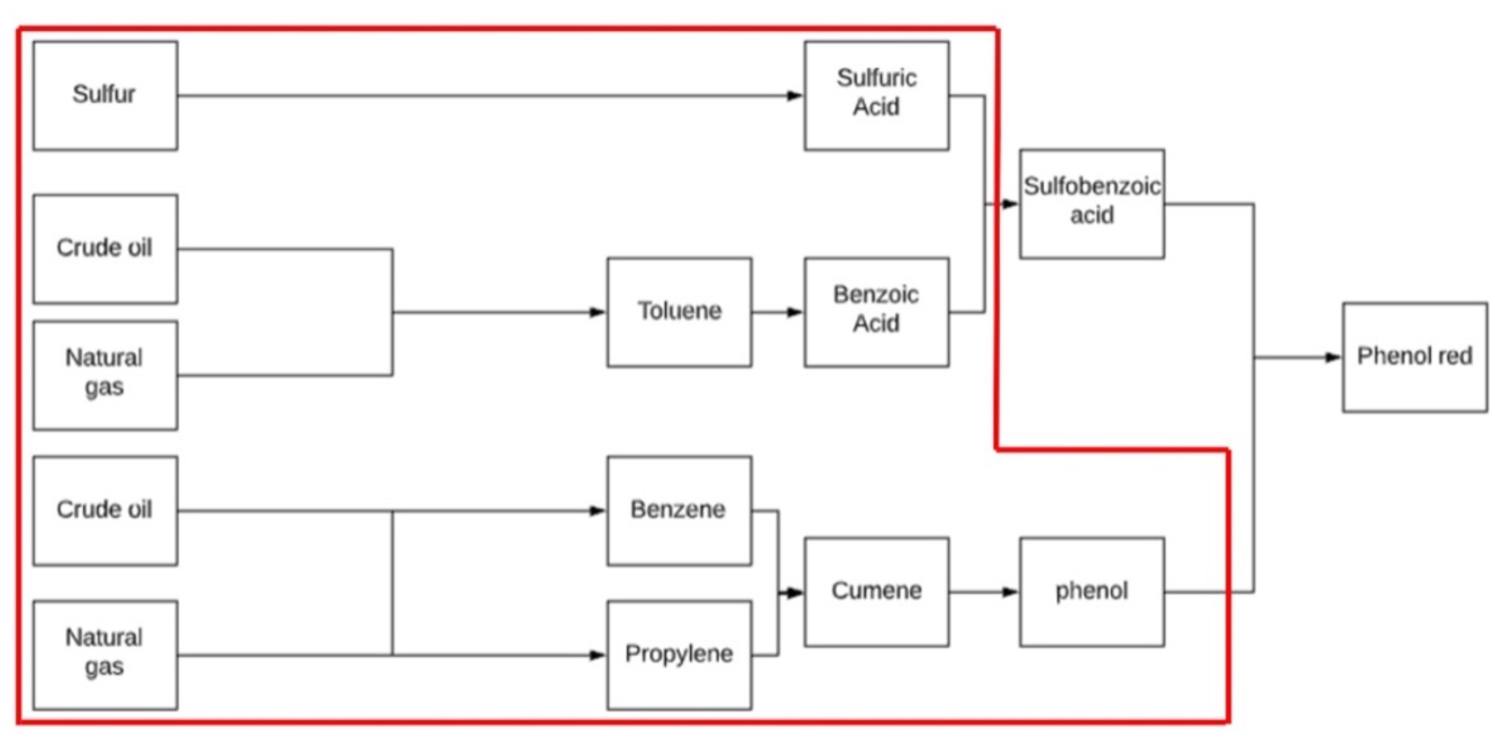
Phenol Red example for ecoinvent dataset utilization. Compounds within the red outline have ecoinvent datasets which account for their material and energy flows. The material and energy flows are not accounted for components outside the red outline. Stoichiometric calculations (theoretical yield utilized) were conducted to estimate mass of each compound when information was not available in ecoinvent (Ex. A minimum of 0.519 kg of sulfobenzoic acid and 0.5311 kg of phenol is needed to produce 1 kg of phenol red).

The material and energy flows for these compounds were tracked and aggregated using OpenLCA software. Table A3 in appendix D provides a list of each component and the components’ precursors. If industrial production information was unavailable, embedded resources could not be quantified, and/or a reasonable substitute could not be identified, then no data were entered for these components. Components without data were still entered into OpenLCA, but without any inputs or outputs. It should be noted that the described method for estimating the inputs and outputs should be considered non-exhaustive due to these gaps in the data.

#### 2.2.5 Solvay and Potash

These categories of E8/B9 components utilize soda ash or potash as major components in their manufacture. For these components, both ecoinvent and available literature estimates were utilized in the same manner as previously described in the chemical category.

#### 2.2.6 Brine evaporation

Sodium chloride is assumed to be produced from a mix of brine and mining operations. Sodium chloride in brine is utilized for soda ash production and is accounted for by utilizing the brine production dataset which does not include cleaning and drying steps. Sodium chloride utilized for other component production processes or as an E8/B9 component are assumed to be produced from a mix of brine and mining operations. The reported embedded resources for non-soda ash related sodium chloride production include extraction, drying, and purification.

#### 2.2.7 Animal cell-produced product

TGF-b can be produced using animal cell culture (Beatson et al., 2011; Zou & Sun, 2004). One advantage of producing TGF-b via animal cell culture rather than a more traditional fermentation organism like *Escherichia coli* is absence of endotoxin. One disadvantage is that the growth medium must be suitable for animal cell culture which has more complex nutrient requirements. This analysis assumes that TGF-b was produced via animal cell culture. Chinese hamster ovary (CHO) cells are the most used animal cell line and are particularly important for glycoprotein overexpression (Beatson et al., 2011; Zou & Sun, 2004). CHO cells require a more complex growth medium as compared to more basic media inputs used for bacteria or yeast growth. DMEM/F12 was utilized as the basal medium for E8/B9 and was deemed to be an acceptable growth medium for CHO cells. The CHO cells were assumed to not require the other 7 components of E8/B9 (ascorbic acid 2-phosphate, additional NaHCO_2_, sodium selenite, insulin Transferrin‡ and FGF-2) (L. Specht, 2019). The material and energy flows were estimated for TGF-b utilizing the data collected for the basal medium production and reported titers of TGF-b.

#### 2.2.8 E8/B9 Components and precursors utilized in multiple production processes

Several E8/B9 component precursors are utilized in the production of multiple E8/B9 components. The material and energy flows necessary to produce these components were accounted for utilizing ecoinvent datasets (ecoinvent Association, 2021). Appendix G lists the components that are utilized in the production of multiple E8/B9 components.

#### 2.2.9 Components not included in assessment

The following components are not accounted for due to the authors’ inability to find either production data or environmental impact data.

##### 2.2.9.1 Lipoic Acid

Lipoic acid was first chemically synthesized in the 1950s (Colingsworth et al., 1952). At an industrial scale, lipoic acid is currently chemically synthesized in three stages (National Center for Biotechnology, n.d.). There has been interest expressed in utilizing biotechnology for lipoic acid production and the overproduction of lipoic acid has been reported in genetically modified *E. coli* (Sun et al., 2017). Despite the advances in biotechnology, lipoic acid is largely produced via chemical synthesis. Data related to the energy and material flows for the chemical synthesis of lipoic acid were not available at time of publication and thus are not included in the LCA.

##### 2.2.9.2 HEPES

A-2-Hydroxyethylpiperazine-N’-2-ethanesulfonic acid, HEPES is a hydrogen ion buffer which is commonly utilized in cell culture (Good et al., 1966). The procedure for HEPES production was first described in 1966 (Good et al., 1966). Data related to industrial manufacture of HEPES was not able to be obtained at time of publication, so the embedded resources associated with HEPES production were not included in the LCA.

#### 2.2.10 Additional B9 components

The composition of B9 is similar to E8, but has additional components: neuregulin, antibiotics/antimycotic, ultrapure water and recombinant albumin. Additional analysis was conducted to evaluate the environmental impact of these supplemental components. antibiotic/antimycotic production typically utilizes 100 kg of solvent and 50 kg of water per kilogram of compound produced (Ho et al., 2010). An ecoinvent-provided equal mix of 15 different organic solvents (acetone, butanol, cumene, cyclohexanol, dichloromethane, ethyl benzene, ethyl glycol, isopropanol, methanol, methyl ethyl ketone, nitrobenzene, styrene, tetrachloroethylene, toluene and xylene) was utilized to estimate the impact of generic organic solvent use. The neuregulin and recombinant albumin environmental impacts were estimated utilizing reported titers (5 mg/L and 17 g/L, respectively) and the method described in microbial-titer methods section (Mautino et al., 2004; Zhu et al., 2021).

#### 2.2.11 Transportation resources

The ecoinvent database v.3.8 was utilized to provide an estimate of the transportation-related resources for each E8/B9 component and their precursors when available. When available transportation was accounted for utilizing ecoinvent datasets which estimate transportation requirements for each product. The ecoinvent datasets provide data on metric ton-km which can be converted to energy via the energy intensities of different modes of transport (MJ/metric ton-km). The energy intensities can vary depending on location and type of transport (Fraser et al., 1995; Gucwa & Schäfer, 2013). OpenLCA software was utilized to consolidate transportation requirements of all inputs and estimate the combined environmental impact of transportation. Appendix H has additional information related the accounting of resources used for transportation.

### 2.3 Life cycle impact assessment

The life cycle impact assessment (LCIA) was conducted utilizing the OpenLCA program v.1.10 and OpenLCA LCIA v2.1.2 methods software. The total direct and indirect energy used throughout the lifecycle of a product, known as, cumulative energy demand (CED) and the tool for reduction and assessment of chemicals and other environmental impacts (TRACI) 2.1 were utilized as the LCIA methods in OpenLCA. CED provides a broad understanding of the energy intensity of processing operations, and also allows for the separation of the estimated energy demand by energy source and renewable or non-renewable classification (Huijbregts et al., 2006). TRACI is an often-cited LCIA tool which utilizes peer-reviewed characterization factors to provide metrics for ozone depletion, climate change, acidification, eutrophication, smog formation, human health impacts, and ecotoxicity (J. Bare, 2011; J. C. Bare et al., 2003). The use of the established CED and TRACI LCIA frameworks provides environmental impact metrics which are reproducible and standardized so that the results are comparable with other product systems. Sensitivity and scenario analyses were also conducted to examine potential uncertainty in the LCIA results and to explore the additional environmental impacts associated with the production of high purity products.

### 2.4 Sensitivity analysis and scenario analysis

A “one-at-a-time” sensitivity analysis was conducted to examine how increases in concentration of each E8/B9 component affect the environmental impact of the functional unit (1 liter E8/B9). It was found that the basal medium, i.e. the core component for both E8 and B9 growth mediums, was responsible for >90% of the LCIA outputs, so an additional one-at-a-time analysis was conducted on the core components of the basal medium. The concentration of each basal medium component was individually increased by 25% and the LCIA calculations for cumulative energy demand and TRACI 2.1 were conducted and recorded using OpenLCA. The percentage of change from the original values was used as an indicator of sensitivity for each variable. See appendix I for sensitivity analysis results.

While LCA can be an important decision-making tool for stakeholders, there is a significant level of uncertainty in the results (Igos et al., 2019). This uncertainty can arise due to a variety of reasons, including but not limited to: generalizations, estimations, spatial considerations, model assumptions, and limited availability of information. To address the uncertainty in our assessment, we conducted a scenario analysis on the E8 growth medium to gain an additional understanding of the potential environmental impacts of producing a highly refined growth medium capable of animal cell culture at currently reported cell densities (cells/ml). We believe the results of the scenario analysis are highly transferable to B9 given its similar composition to E8. We examined three scenarios as described below.

#### Baseline scenario

This scenario accounts only for the data which we were able to obtain from our described methods. It does not account for any additional processing or resources which are associated with pharmaceutical grade ingredient production. This scenario should be considered a minimum due to the limited nature of the analysis which is described in the method section.

#### Partial purification scenario

This scenario increases the environmental impact of non-basal media components of E8 (ascorbic acid 2-phosphate, additional NaHCO_3_, sodium selenite, insulin, transferrin and FGF-2) by 20-fold to account for additional processing associated with active pharmaceutical ingredient production (Wernet et al., 2010). The increase in impact accounts for the additional energy and resources used for the purification process.

#### High purification scenario

This scenario increases the environmental impact of all components by 20-fold to examine the impact of all E8 components being processed to the purity of active pharmaceutical products (Wernet et al., 2010). Again, the increase in environmental impacts derives from the additional energy and resources used for the purification process.

## 3. Results

The baseline results indicate a dramatic difference in E8 and B9, however this difference can be attributed to the inclusion of antibiotics in the B9 formulation. When an antibiotic-free version of B9 (B9af) is considered, the energy use and environmental impacts are analogous to E8. We examined the LCIA results for CED and TRACI LCIA methods for the E8, B9 and B9af.

OpenLCA attributed the majority of the environmental impacts (>90%) of both E8 and B9af to the DMEM/F12 basal medium. To further analyze the environmental impacts of DMEM/F12, a sensitivity analysis was conducted on each DMEM/F12 component. This analysis found that glucose was the most environmentally impactful component of the DMEM/F12 medium and this is largely due to its relatively high concentration (3.151g/L) in relation to the other DMEM/F12 growth medium components. Additionally, our scenario analysis of the E8 growth medium highlights the significant influence of high levels of refinement/purification of the E8 components on increasing the environmental impact of the growth medium production.

For E8, the DMEM/F12 basal medium was the most environmentally impactful constituent when considered as a single ingredient/component of the media composition. Figure 3 provides a breakdown of the energy sources used for growth medium production and provides the total CED for a liter of each growth medium. The change in the total CED between the DMEM/F12 basal medium and E8 was ∼4%. This change is even less significant when comparing DMEM/F12 growth medium and the B9af growth mediums (Figure 3.0). The replacement of a portion of DMEM/F12 with ultrapure water in B9af growth medium can be attributed with the lower CED of B9af. However, when the addition of antibiotics/antimycotic is accounted for in B9 the total CED increases nearly two orders of magnitude when compared to E8 or B9af (390 MJ/L vs. ∼1.7 MJ/L, respectively). This increase can be attributed to the high volumes of organic solvent and water (100 kg of solvent/1 kg of antibiotic/antimycotic and 50 kg of water/1 kg of antibiotic/antimycotic) associated with antibiotic/antimycotic or high-purity small molecule production (Ho et al., 2010). These organic solvents originate from fossil fuels which accounts for the order of magnitude increase in fossil fuel CED.

**Figure 3.0.**
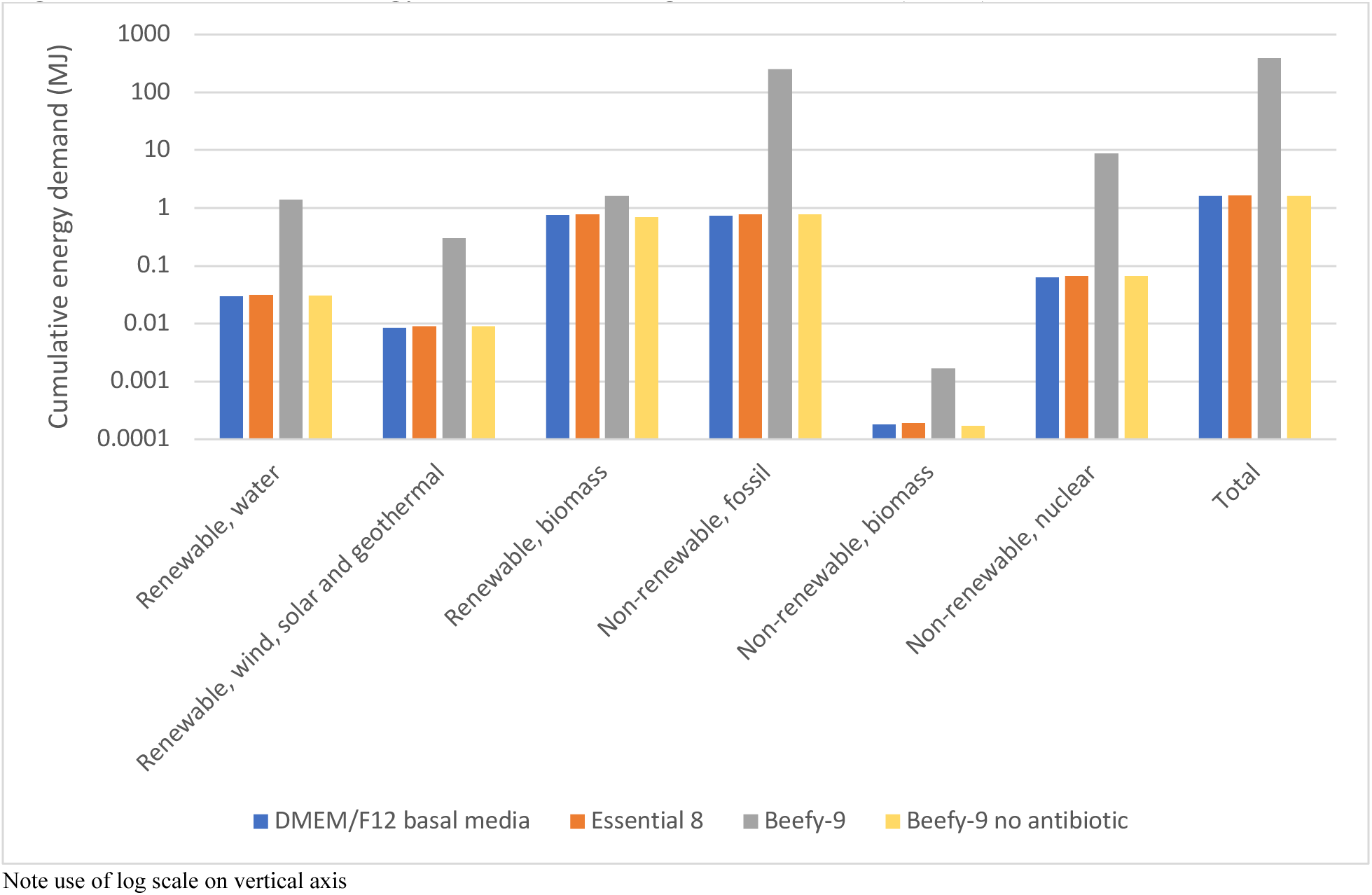
Cumulative energy demand of each growth medium (MJ/L)

The results of the TRACI LCIA indicate minimal differences in E8, DMEM/F12 and B9af growth mediums (Table 2.0). When antibiotic containing growth mediums are included, the B9 TRACI LCIA results are orders of magnitudes higher than E8 and DMEM/F12 growth mediums across most impact categories. For example, the global warming potential (GWP) is ∼122x higher for B9 than E8, and B9 would deplete ∼448x more fossil fuel than E8. Figure 4.0 illustrates the magnitude of change between B9 and B9af. Thus, from an environmental perspective, the reduction and/or elimination of antibiotic/antimycotic growth medium components would be particularly advantageous. It is also important to note that this analysis does not account for the antibiotics being released into the environment during production. Additional analysis would be necessary if an antibiotic containing growth medium is used for industrial scale production of non-vital products.

**Figure 4.0.**
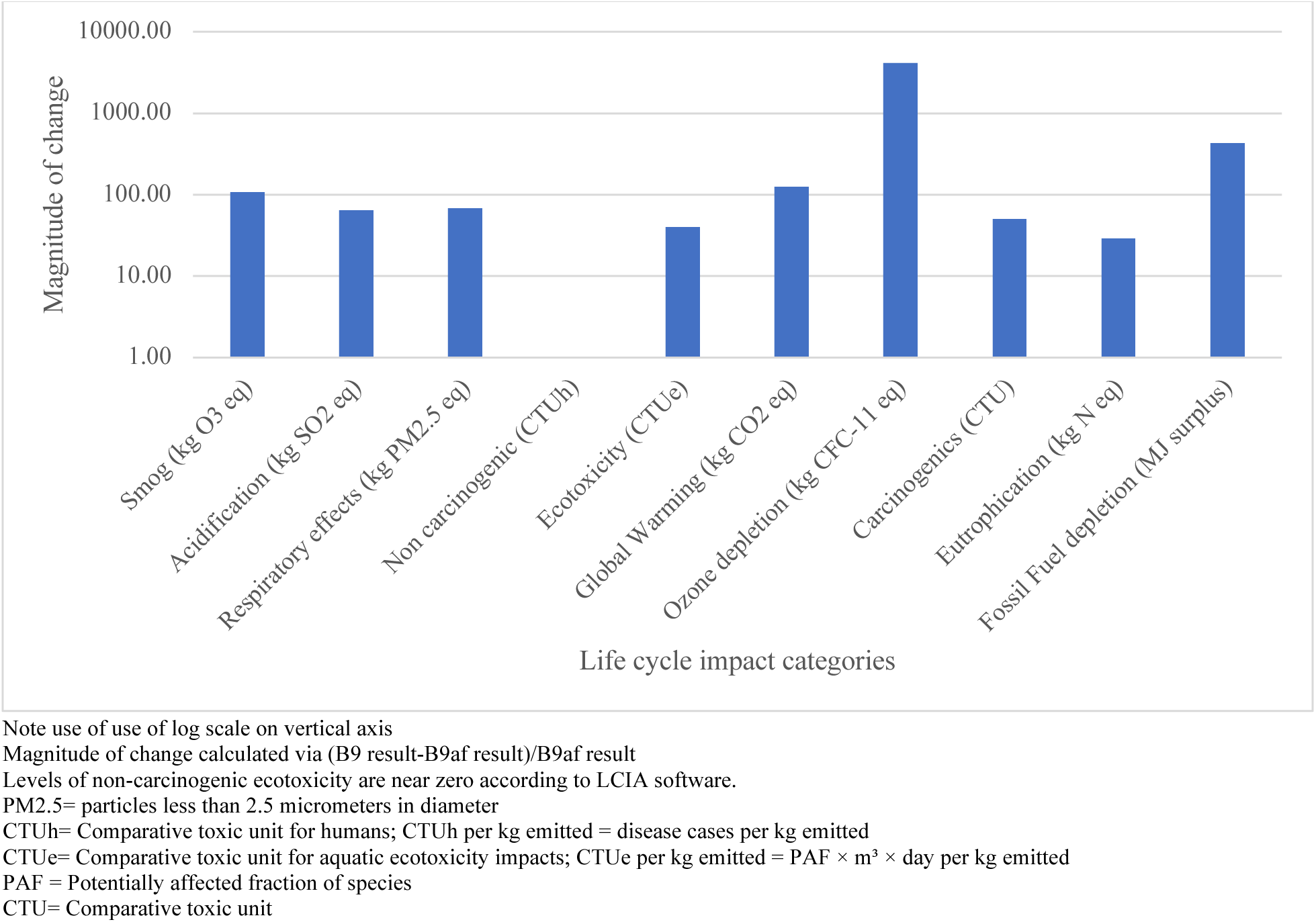
Magnitude of change of each TRACI impact category for B9 relative to B9af

**Table 2.0.** TRACI impact category results for one liter of growth medium

|  | DMEM/F12 basal media | Essential 8 | Beefy-9 | Beefy-9 no antibiotic |
| --- | --- | --- | --- | --- |
| Smog (kg O<sub>3</sub> eq) | 3.66E-03 | 3.89E-03 | 3.73E-03 | 4.06E-01 |
| Acidification (kg SO<sub>2</sub> eq) | 5.30E-04 | 5.60E-04 | 5.20E-04 | 3.43E-02 |
| Respiratory effects (kg PM2.5 eq) | 6.62E-05 | 7.05E-05 | 6.65E-05 | 4.65E-03 |
| Non carcinogenic (CTUh) | -1.62E-08 | -1.56E-08 | -1.36E-08 | 1.08E-06 |
| Ecotoxicity (CTUe) | 1.50E+00 | 1.61E+00 | 1.50E+00 | 6.15E+01 |
| Global Warming (kg CO<sub>2</sub> eq) | 6.20E-02 | 6.57E-02 | 6.40E-02 | 8.03E+00 |
| Ozone depletion (kg CFC-11 eq) | 5.75E-09 | 6.00E-09 | 7.11E-09 | 2.92E-05 |
| Carcinogenics (CTU) | 7.09E-09 | 7.55E-09 | 7.07E-09 | 3.65E-07 |
| Eutrophication (kg N eq) | 3.80E-04 | 3.90E-04 | 3.90E-04 | 1.18E-02 |
| Fossil Fuel depletion (MJ surplus) | 7.10E-02 | 7.43E-02 | 7.70E-02 | 3.33E+01 |

\*Levels of non-carcinogenic ecotoxicity reported as near zero negative and positive values according to LCIA software
PM2.5= particles less than 2.5 micrometers in diameter
CTUh= Comparative toxic unit for humans; CTUh per kg emitted = disease cases per kg emitted
CTUe= Comparative toxic unit for aquatic ecotoxicity impacts; CTUe per kg emitted = PAF × m<sup>3</sup> × day per kg emitted
PAF = Potentially affected fraction of species
CTU= Comparative toxic unit

A scenario analysis was conducted to further explore the potential environmental impacts of media production, including the additional impacts of producing high-purity or pharmaceutical grade E8 components (Wernet et al., 2010). The scenario analysis examined the most established growth medium, E8, and was utilized to examine how the environmental impact might change as the level of refinement increases for some or all E8 components. The first scenario is the baseline scenario and should be considered the minimum environmental impact of E8 production due to the limited nature of this study. The CED for the baseline scenario was 1.65 MJ/per liter of E8 with the majority of the energy being supplied by non-renewable fossil fuel and renewable biomass. This is greater than seven times the amount of energy used to light a 60-watt incandescent lightbulb for an hour.

Partial purification scenario increased the seven components other than DMEM/F12 growth medium by a factor of 20 to account for the additional impact associated with high purity/pharmaceutical compound production (Wernet et al., 2010). This increase in partial purification scenario nearly doubled the total CED compared to the first scenario. This highlights the potential impact that high purity substances can have on total CED for the production process.

The final scenario examines the potential environmental impacts if all E8 components (including DMEM/F12) are produced as high purity compounds with their potential resource use being increased by a factor of 20. The total cumulative energy demand for scenario 3 was 33 MJ/L of E8, which is more energy that is contained in a liter of gasoline (Engineering Toolbox, 2008). This indicates that the level of purification and refinement of each component will heavily influence the cumulative energy demand of E8 (Figure 5.0). Orders of magnitude differences in the TRACI 2.1 outputs can be observed within the scenarios as well.

**Figure 5.0.**
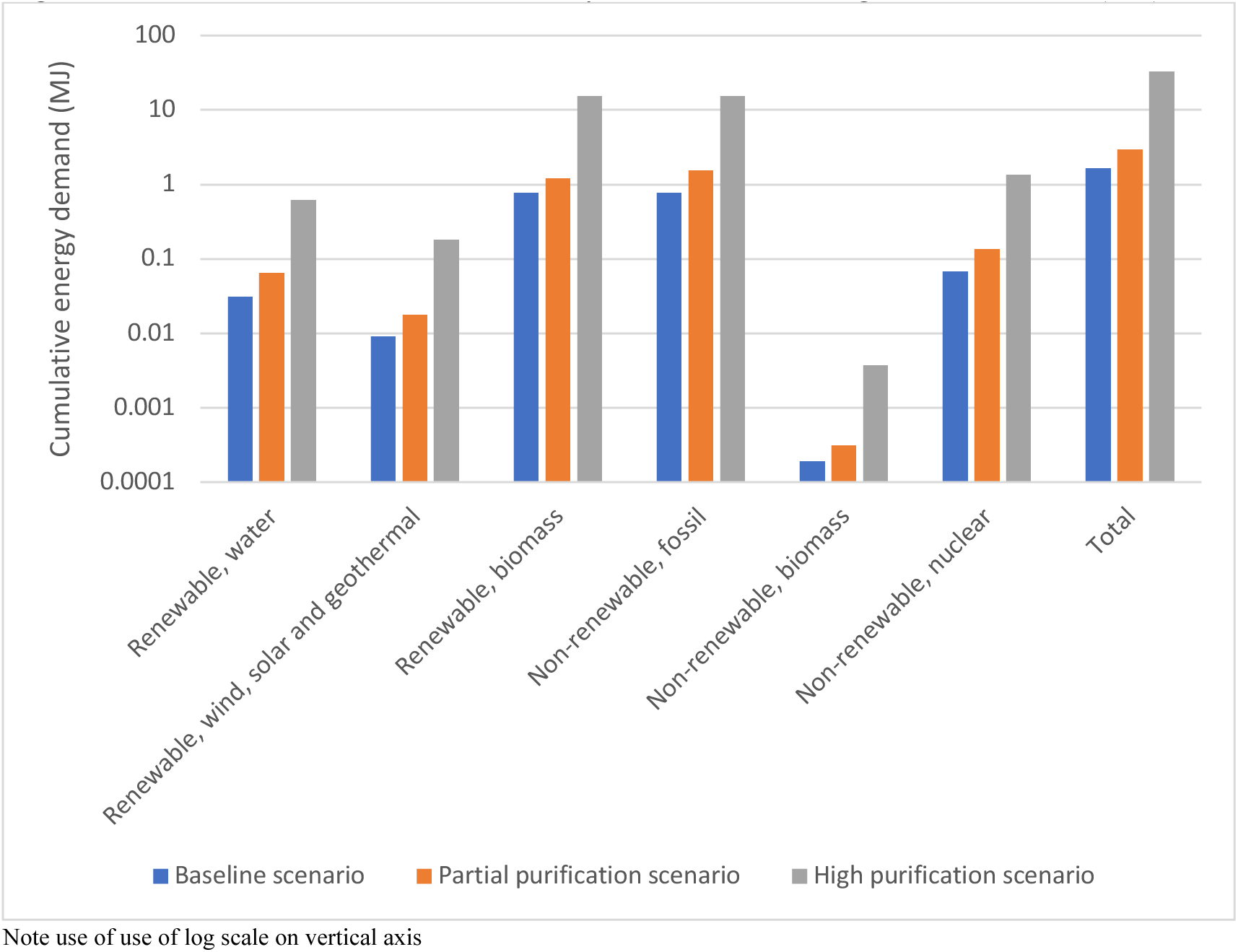
CED results for scenario analysis of Essential 8 growth medium (MJ)

The TRACI 2.1 results for each scenario are similar to the order of magnitude changes observed in the cumulative energy demand. The GWP increases linearly from the baseline scenario to highly purified scenario, reflecting the 20x increase associated with complete reliance on high purity inputs for Scenario 3. The GWP doubles between the baseline scenarios and partial purification scenario which indicates that utilizing a few high purity E8 components can substantially increase the environmental impact of the E8 production. Also, it should be recognized that the baseline scenario is the minimum GWP of E8 production. The fossil fuel depletion impact category increased by 20x when comparing the baseline scenario and high purification scenario. The overall TRACI 2.1 results indicated that a ∼20x increase in resource use produced roughly linear results in each impact.

If we examine the top three ecoinvent datasets for each impact category it becomes apparent certain datasets have the largest impact on each TRACI 2.1 impact category (see Figure 6.0). It appears that glucose production, glucose transport (market for…) and maize transport have outsize impacts in multiple categories. This is likely due to the reported concentration of glucose (∼3 g/L) in the basal medium with only NaCl and HEPES being reported to having a higher concentration (∼7 g/L and 3.5 g/L, respectively). It should again be noted that the environmental impacts of HEPES buffering agent production were not accounted for due to the author’s inability to obtain industrial production data.

**Figure 6.0.**
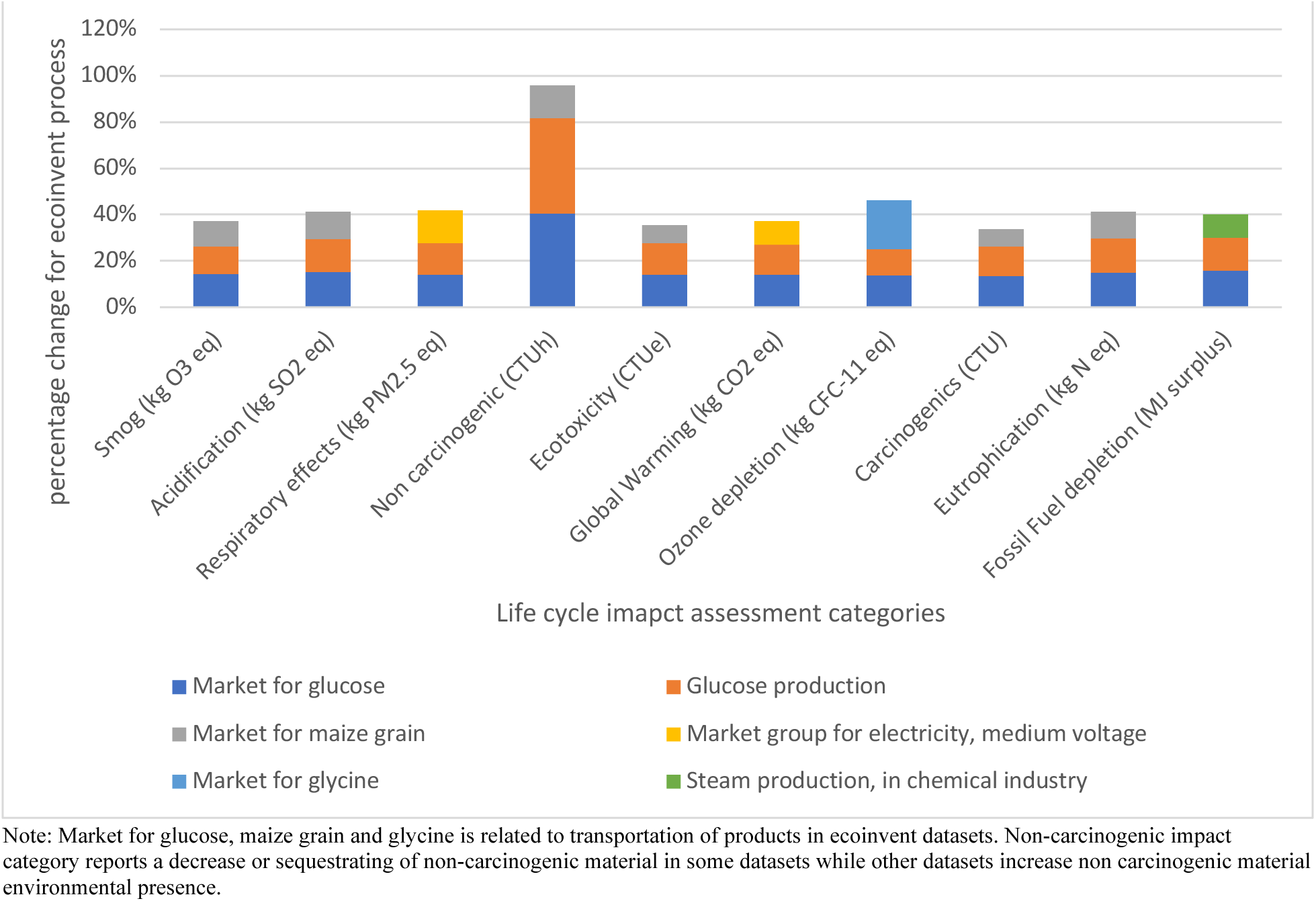
Absolute percentage change in each of the top three ecoinvent datasets for each TRACI 2.1 impact category

## 4. Discussion

Understanding the minimum environmental impacts of animal/human cell growth media provides a starting point for developing an in-depth understanding of the environmental impact of animal cell-based bioproducts. This LCA provides existing biomanufacturers and laboratories with data that can be utilized to help better understand the current environmental impacts of their operations. The data from this LCA can also be used to provide environmental impact metrics for increases in production efficiency (increased titer) or reduction in overall growth medium usage. The economic importance of increasing a production facility’s titer, production scale, or reducing growth medium use is often cited (Humbird, 2021; Risner et al., 2020; Wu & Maravelias, 2018), however our LCA provides the metrics to quantify the potential positive environmental impacts of any advancements to improve the efficiency of growth medium use.

Understanding the potential environmental impact of E8/B9 consumption can also help with the environmentally responsible scaling of nascent biotechnology industries, such as the animal cell-based meat or stem cell therapy industries. Researchers and companies currently use highly refined growth mediums for animal cell culture and understanding the environmental ramifications of their work may be important to their environmentally and socially conscious investors or financiers. This consideration may be less important for manufacturers of therapeutics, but it should be an important consideration for those seeking to change large-scale commodity systems like our food and agriculture systems with environmental goals in mind.

These emerging industries could use this initial study to understand the near-term environmental implications of developing large scale systems before maximizing metrics like achievable cell concentration (cells/ml). Advances in technology which reduce growth medium use and/or increase the achievable cell concentration will play an important economic and environmental role for biobased products particularly those which seek to be alternatives to commodity-type products, such as animal cell-based meat. This convergence of economic and environmental interest is one of the strengths of the developing bioeconomy, however this work also highlights the potential environmental challenges of utilizing large volumes of growth medium for bioproduct production.

The reported environmental impacts of the baseline scenario should be considered the minimum environmental impacts due to some key limitations in the analysis. First, the overall system boundary was partially truncated since we were unable to fully account for all E8/B9 components such as HEPES and lipoic acid. Incomplete LCI information in ecoinvent and the literature (See Methods section) also contributed to the limited assessment of the environmental impact of some components in the growth medium. It should be noted that the HEPES buffering agent whose concentration was greater than glucose (3.575 vs. 3.151g/L) was not accounted for and may substantially contribute to the environmental impact given the concentration and its chemical synthesis. The environmental impacts of other compounds, such as phenol red were only partially accounted for as well. In addition, the assumption of utilizing microbial fermentation to produce some compounds may not be true in all cases. This highlights why the reported environmental impacts of this LCA should be considered as a minimum.

E8 is an established animal cell growth medium which has been used for animal cell research since at least 2011 and does not include antibiotics within its formulation (Chen et al., 2011). B9 which includes antibiotics/antimycotics and other additional components (neuregulin, ultrapure water, and recombinant albumin) has yet to be commercially established (Stout et al., 2021). If antibiotic use is required for B9 to be a viable growth medium, then its environmental impact is likely to be two orders of magnitude greater than E8. If the B9 growth medium can be utilized without the addition of antibiotics, then our findings indicate that these growth mediums have highly similar environmental impacts. This indicates that from an environmental standpoint, antibiotic use should be limited/eliminated from any large-scale cell proliferation system. It is also worth noting that this LCA also does not evaluate the complex potential health and environmental implications of antibiotics entering our water or terrestrial systems (J. Wang et al., 2020).

When E8 and B9af were compared it was found that DMEM/F12 basal medium was the major contributor to the environmental impacts of both growth mediums. A sensitivity analysis was conducted which analyzed the environmental impact of the components of DMEM/F12 basal medium. The sensitivity analysis found that glucose was the most impactful component and approximately half of glucose’s global warming potential can be attributed to its transport (26%) and production from starch slurry (24%). However, due to the quantity of DMEM/F12 basal medium components (>50), there was no single component that could be attributed to the majority (>50%) of the environmental impacts. Approximately half of glucose’s global warming potential can be attributed to it’s transport (26%) and production from starch slurry (24%). Our results should be transferable to growth mediums which utilize DMEM/F12 as a basal medium. However, additional analysis would be needed for growth mediums whose basal medium composition are different than DMEM/F12’s composition.

The scenario analysis explores how increased levels of purification which are required for pharmaceutical or fine chemical production can increase the embedded resources and energy within a growth medium. The results of the partial and high purification highlight how the potential refinement of some or all growth medium components can influence the environmental impacts of a growth medium like E8. Additional energy production and use is responsible for 65-85% of the additional environmental impacts associated with fine chemical production (Wernet et al., 2010). This indicates that employing energy efficient means of refinement will be important for the sustainable production of growth medium components, and especially critical for applications which may require high volumes of growth medium. All components may not require the same level of refinement as fine chemicals but utilizing less refined components has the potential to increase the risk of contamination, batch variation, or failure.

An additional challenge related to cell culture purification is endotoxin removal. Endotoxin is a heat and pH stable lipopolysaccharide with a molar mass ranging from 2.5-70 kDa, and if it is introduced to the bioreactor via the growth medium, it has the potential to limit cell proliferation (Magalhães et al., 2007). These characteristics can make endotoxin removal challenging and different purification strategies must be deployed based on the substance’s properties (EMD Millipore, 2012). The heat stability of endotoxin can make traditional steam sterilization protocols (121°C for 45 min or 132°C for 4 min) ineffective since inactivation requires temperatures 250 °C for 30 min or 180 °C for 3 h (Centers for Disease Control and Prevention, 2016; Magalhães et al., 2007). Other methods of removal which utilize filtration or charged/hydrophobic membrane interactions are highly dependent upon the characteristics of the proteins/compounds which are being purified (EMD Millipore, 2012). A variety of these methods have been utilized to separate and remove endotoxins from laboratory grade components including LPS affinity resins, two-phase extractions, ultrafiltration, hydrophobic interaction chromatography, ion exchange chromatography, and membrane adsorbers (Magalhães et al., 2007). Of course, these additional processing steps would likely increase production costs as well as the environmental impact of the E8/B9 production. Due to these purification challenges, it is likely worth prioritizing the development of an efficient and environmentally friendly method of endotoxin removal to reduce the environmental impact of E8/B9 production.

## 5. Conclusion

The bioeconomy has been touted as one of the solutions to the United Nations SDGs, however critical environmental, economic and social assessment is needed to understand the sustainability of biobased products. An environmental assessment of animal or human stem cell growth mediums (E8 and B9) is necessary to understand the environmental impact of a multitude of potential products ranging from therapeutic stem cell therapy to ACBM products. Further, individually specified sustainability assessments of biobased products are necessary due to the multitude of different factors (e.g. type of production organism, doubling time, product yield, and titer, among other factors) that influence resource use and environmental impact.

This LCA provides a foundational understanding of the minimum, near-term environmental impacts of growth mediums utilized for animal cell proliferation. It examines the environmental impacts of an established stem cell growth medium (E8) and compares it with an emerging growth medium, B9. It was found that antibiotic/antimycotic use is highly environmentally impactful, however if an antibiotic free version of B9 (B9af) is utilized then the E8 and B9af environmental impacts are similar. The similarity in environmental impacts of E8 and B9af can be attributed to the fact both mediums utilize DMEM/F12 as the basal medium. Scenarios are utilized to explore how increased levels of refinement and purification of growth medium components can potentially increase the environmental impact of a growth medium. The environmental impacts in this LCA should be viewed as the minimum impacts of E8, B9, B9af and DMEM/F12 production due to the truncated nature of this assessment. This LCA provides a starting point for researchers working at the convergence of emerging animal cell-based biotechnology and sustainability.

The quantified environmental impacts of these human and animal cell growth mediums will be an essential resource for assessing the environmental impacts of emerging bioproducts. Further, this information can be used to understand how the reduction in growth medium use or an increase in titer can have a positive environmental impact. This work acts as a foundation for future LCAs or other environmental assessments which examine products produced from animal or human stem cells. Future work could involve a more regional assessment of each growth medium component, detailed assessment of individual growth medium components, impact assessment of missing E8/B9 components (Ex. HEPES), the development of environmentally friendly bioprocesses and/or sustainability assessments of near-term human or animal cell products.

## 6. Appendices

### Appendix A- Raw ingredients

#### Glucose

The material flow for glucose production was utilizing ecoinvent 3.8 datasets. The maize was assumed to be dried during initial harvest and flow the can be described in calculation A1. The conversion factors were all taken from the ecoinvent 3.8 datasets. Transportation was assumed between each step and is accounted for utilizing ecoinvents datasets.

Calculation A1. Example of ecoinvent mass determination: Maize mass needed for glucose production

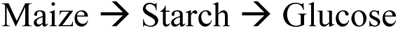

1 kg of glucose x 0.9 kg starch/1 kg glucose = 0.9 kg of starch

0.9 kg of starch x 1.261 kg of dry corn/1 kg of starch = 1.1349 kg of dry corn

Calculation notes:

1 kg of glucose requires 0.9 kg of starch to produce (ecoinvent Association, 2021)

1 kg of starch requires 1.261 kg of maize (ecoinvent Association, 2021)

#### Linoleic acid

Cottonseed oil production was utilized to estimate the linoleic acid production. The separation of linoleic acid from fatty acids was not accounted for and would likely increase the input/outputs of this E8/B9 component. The ecoinvent datasets were utilized to estimate the material/energy flow from cottonseed production to cottonseed oil refining. Transportation was assumed between each production step. Refined cottonseed oil was assumed to have a similar triglyceride content (96% w/w) (Matthäus, 2010). Cottonseed oil’s fatty acid profile has been reported consist of the 52% linoleic acid. This indicates that a minimum of ∼2 kg of refined cottonseed oil would be required to produce 1 kg of linoleic acid.

### Appendix B- Microbial yield

**Table A1.** Reported microbial yields and calculated correction factor for some E8/B9 components

| Compound | Yield<br>(g/g<br>glucose) | Correction<br>factor | Source |
| --- | --- | --- | --- |
| i-Inositol <sup>1</sup> | 0.820 | 0.788 | (Y. Li et al., 2021) |
| L-Alanine | 0.520 | 1.24 | (Drauz et al., 2007) |
| L-Arginine-HCl <sup>2</sup> | 0.431 | 1.50 | (Man et al., 2016) |
| L-Asparagine-H<sub>2</sub>O <sup>2</sup> | 0.836 | 0.773 | (Drauz et al., 2007) |
| L-Cysteine-HCl-H<sub>2</sub>O <sup>2</sup> | 0.060 | 10.8 | (H. Liu et al., 2018) |
| L-Cystine <sup>2</sup> | 0.060 | 10.8 | (H. Liu et al., 2018) |
| L-Glutamic acid | 0.630 | 1.03 | (Wen & Bao, 2019) |
| L-Histidine-HCl-H<sub>2</sub>O <sup>1,2</sup> | 0.080 | 8.08 | (Schwentner et al., 2019) |
| L-Isoleucine | 0.300 | 2.15 | (Drauz et al., 2007) |
| L-Leucine | 0.150 | 4.31 | (Drauz et al., 2007) |
| L-Lysine-HCl <sup>2</sup> | 0.646 | 1.00 | (Félix et al., 2019) |
| L-Methionine<sup>1</sup> | 0.130 | 4.97 | (Zhou et al., 2019) |
| L-Phenylalanine | 0.180 | 3.59 | (Drauz et al., 2007) |
| L-Proline | 0.360 | 1.79 | (Wendisch et al., 2016) |
| L-Serine | 0.430 | 1.50 | (Zhang et al., 2018) |
| L-Threonine | 0.400 | 1.62 | (Drauz et al., 2007) |
| L-Tryosine | 0.100 | 6.46 | (Lütke-Eversloh et al., 2007) |
| L-Tryptophan | 0.200 | 3.23 | (Drauz et al., 2007) |
| L-Valine | 0.280 | 2.31 | (Drauz et al., 2007) |
| Pyridoxal-HCl <sup>2</sup> | 0.325 | 1.99 | (Y. Wang et al., 2021) |
| Pyridoxine-HCl <sup>2</sup> | 0.325 | 1.99 | (Y. Wang et al., 2021) |
| Sodium Pyruvate <sup>2</sup> | 0.678 | 0.953 | (Y. Li et al., 2001; Miyata & Yonehara, 1999) |
| Thymidine | 0.019 | 34.0 | (Lee et al., 2010) |

<sup>1</sup>Coverted from mol/mol<sub>glucose</sub> to g/g<sub>glucose</sub>
<sup>2</sup>See information below for additional clarification
Calculation A2. Correction factor calculation example for compounds with reported yields
\*L-Lysine yield (g/g<sub>glucose</sub>) constant in all calculations
\*\*L-Leucine (g/g<sub>glucose</sub>)

**Table A2.**
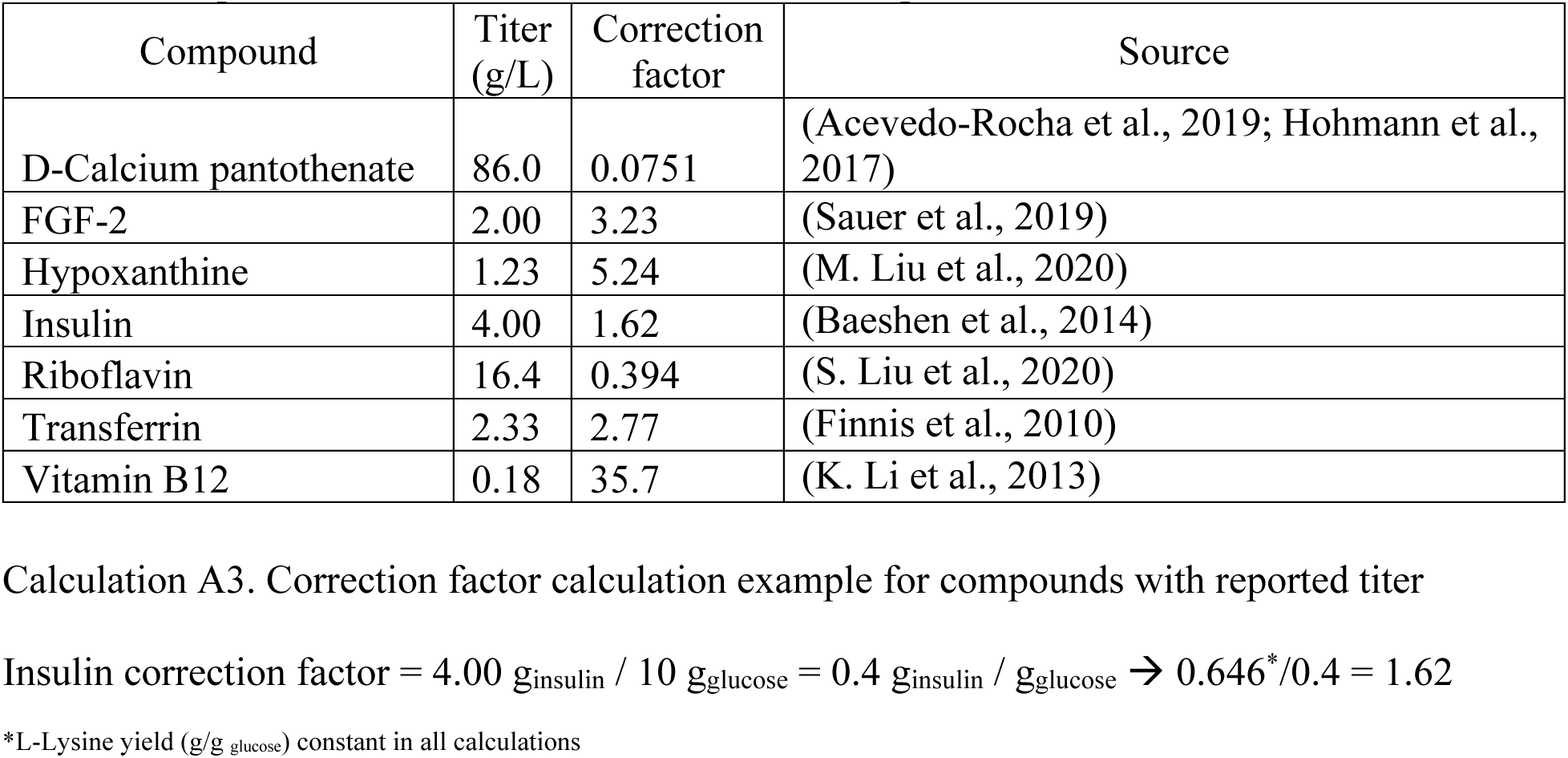
Reported microbial titer for some E8/B9 components

| Compound | Titer (g/L) | Correction factor | Source |
| --- | --- | --- | --- |
| D-Calcium pantothenate | 86.0 | 0.0751 | (Acevedo-Rocha et al., 2019; Hohmann et al., 2017) |
| FGF-2 | 2.00 | 3.23 | (Sauer et al., 2019) |
| Hypoxanthine | 1.23 | 5.24 | (M. Liu et al., 2020) |
| Insulin | 4.00 | 1.62 | (Baeshen et al., 2014) |
| Riboflavin | 16.4 | 0.394 | (S. Liu et al., 2020) |
| Transferrin | 2.33 | 2.77 | (Finnis et al., 2010) |
| Vitamin B12 | 0.18 | 35.7 | (K. Li et al., 2013) |

Calculation A3. Correction factor calculation example for compounds with reported titer
\*L-Lysine yield (g/g glucose) constant in all calculations

#### L-Arginine-HCl

Microbial L-arginine production is utilized as a substitute for L-arginine-HCl production. Additional resources with processing L-arginine into L-arginine-HCl-H_2_O are not accounted for.

#### L-Asparagine-H_2_O

L-Asparagine can be synthesized utilizing L-aspartic acid which is esterified followed by treatment with ammonia (Drauz et al., 2007). To estimate the embedded resources in asparagine production, the yield (.836 g/g glucose) for aspartic acid production was utilized. The resources utilized for esterification, ammonia treatment and additional steps for the conversion of L-asparagine to L-asparagine H_2_0 are not accounted for.

#### L-Cysteine-Cl-H_2_O

Microbial L-cysteine production is utilized as a substitute for L-cysteine-HCl-H_2_O production. Additional resources with processing L-cysteine into L-cysteine-HCl-H_2_O are not accounted for.

#### L-Cystine

It is the oxidized dimer formed from a pair of cysteine molecules. The reported yield for the microbial production of cysteine is 0.06 g/g glucose and this value is utilized for cystine production (H. Liu et al., 2018).

#### L-Histidine-HCl-H_2_O

Microbial L-histidine production is utilized as a substitute for L-histidine-HCl-H_2_O production. Additional resources with processing L-histidine into L-histidine-HCl-H_2_O are not accounted for.

#### Sodium Pyruvate

Microbial pyruvate production is utilized as a substitute for sodium pyruvate production. Additional resources with processing pyruvate into sodium pyruvate are not accounted for.

#### Pyridoxal-HCl and Pyridoxine-HCl

Pyridoxal HCl and pyridoxine HCl are forms of B_6_ and have been produced microbially via recombinant *Sinorhizobium meliloti* (Acevedo-Rocha et al., 2019; Y. Wang et al., 2021). A titer of 1.3 g/L of B_6_ has been report and this titer was used to estimate yield based upon a minimal media containing 4 g of glucose/L (King, 2015; Y. Wang et al., 2021). The estimated yield was utilized to estimate the embedded resources for both pyridoxal HCl and pyridoxine HCl production and additional processing was not accounted for.

### Appendix C- Enzymatic

#### L-Aspartic acid

L-aspartic acid has been described as being produced enzymatically with yields of up to 0.95 (g/g fumaric acid) being reached while utilizing fumaric acid as a feedstock (Appleton & Rosentrater, 2021; Yukawa et al., 2010). Fumaric acid can be produced utilizing glucose as feedstock with a yield of 0.88 g/g glucose (Martin-Dominguez et al., 2018). To estimate embedded resources, the required mass of glucose was determined (∼1.20 g glucose per g of L-aspartic acid produced). The embedded resources associated with the mass of glucose was then attributed to L-aspartic acid production.

### Appendix D- Chemical

The following section provides the methodology utilized to estimate the environmental impact of E8/B9 components. Table A3 provides the E8/B9 precursor components for each chemical classified E8/B9 component and additional details related to the life cycle inventory methodology can be found in this subsequent section.

**Table A3.**
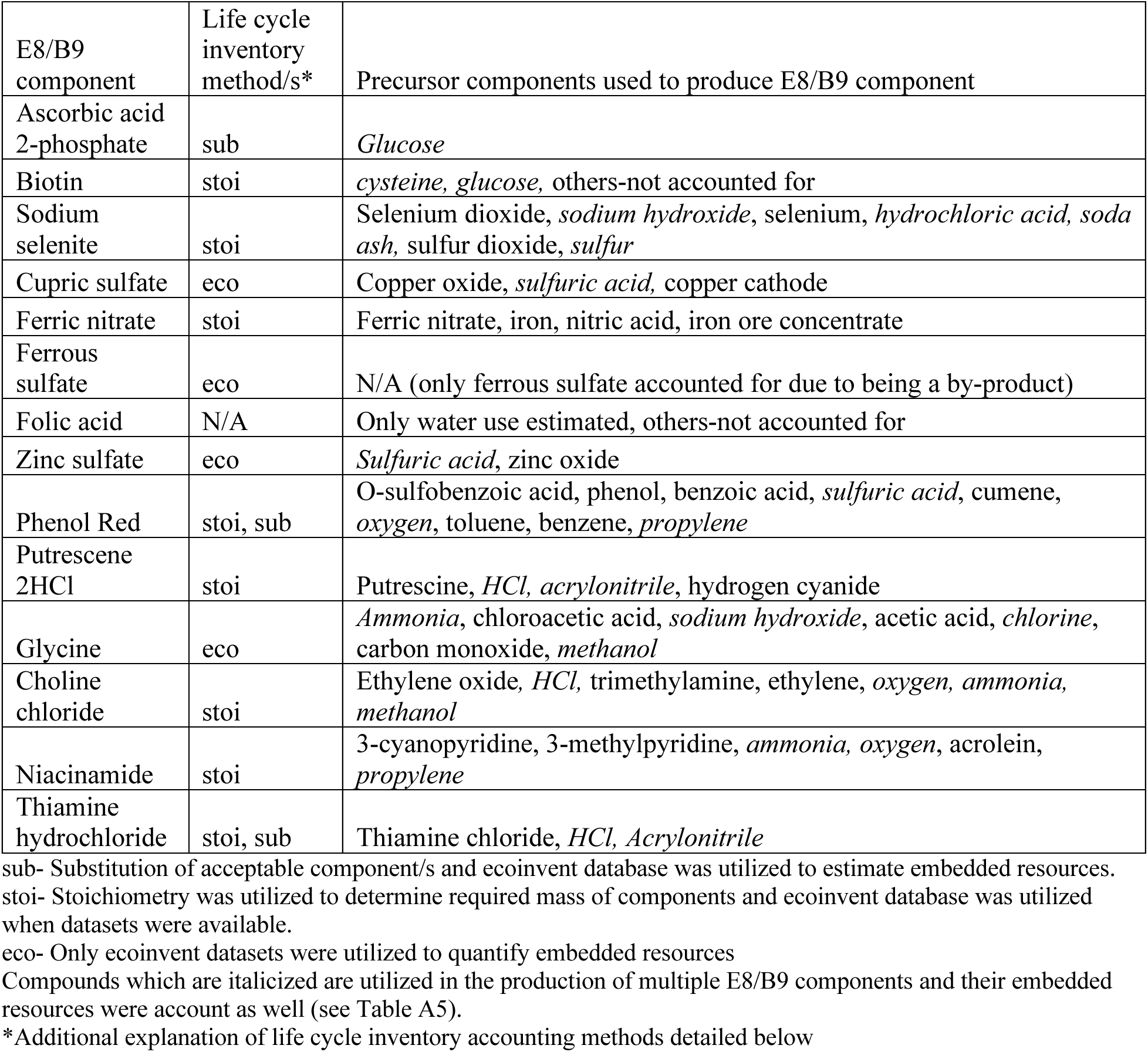
Chemically manufactured E8/B9 components’ life cycle inventory accounting methods and precursors components

| E8/B9 component | Life cycle inventory method/s* | Precursor components used to produce E8/B9 component |
| --- | --- | --- |
| Ascorbic acid 2-phosphate | sub | *Glucose* |
| Biotin | stoi | *cysteine, glucose*, others-not accounted for |
| Sodium selenite | stoi | Selenium dioxide, *sodium hydroxide*, selenium, *hydrochloric acid*, *soda ash*, sulfur dioxide, *sulfur* |
| Cupric sulfate | eco | Copper oxide, *sulfuric acid*, copper cathode |
| Ferric nitrate | stoi | Ferric nitrate, iron, nitric acid, iron ore concentrate |
| Ferrous sulfate | eco | N/A (only ferrous sulfate accounted for due to being a by-product) |
| Folic acid | N/A | Only water use estimated, others-not accounted for |
| Zinc sulfate | eco | *Sulfuric acid*, zinc oxide |
| Phenol Red | stoi, sub | O-sulfobenzoic acid, phenol, benzoic acid, *sulfuric acid*, cumene, *oxygen*, toluene, benzene, *propylene* |
| Putrescene 2HCl | stoi | Putrescine, *HCl*, *acrylonitrile*, hydrogen cyanide |
| Glycine | eco | *Ammonia*, chloroacetic acid, *sodium hydroxide*, acetic acid, *chlorine*, carbon monoxide, *methanol* |
| Choline chloride | stoi | Ethylene oxide, *HCl*, trimethylamine, ethylene, *oxygen*, *ammonia*, *methanol* |
| Niacinamide | stoi | 3-cyanopyridine, 3-methylpyridine, *ammonia*, *oxygen*, acrolein, *propylene* |
| Thiamine hydrochloride | stoi, sub | Thiamine chloride, *HCl*, *Acrylonitrile* |

sub- Substitution of acceptable component/s and ecoinvent database was utilized to estimate embedded resources.
stoi- Stoichiometry was utilized to determine required mass of components and ecoinvent database was utilized when datasets were available.
eco- Only ecoinvent datasets were utilized to quantify embedded resources
Compounds which are italicized are utilized in the production of multiple E8/B9 components and their embedded resources were account as well (see Table A5).
\*Additional explanation of life cycle inventory accounting methods detailed below

#### AA2P (ascorbic acid 2-phosphate)

An ecoinvent dataset for ascorbic acid 2-phosphate was not reported and ascorbic acid was deemed as an acceptable substitute. The required mass of glucose, the major component utilized in ascorbic acid production was determined via E8/B9 concentration of ascorbic acid 2-phosphate and ecoinvent database. The energy and material flows were determined from the ascorbic acid ecoinvent dataset and from the material/energy associated with glucose produced from corn (see raw ingredients: glucose section).

#### Biotin

Biotin can be synthesized from multiple precursors utilizing multi-step reactions (Bonrath et al., 2009; Casutt et al., 2011; de Clercq, 1997; Tang et al., 2020). Multiple reaction schemes have been reported to be used for the conversion of cysteine to biotin and we assume cysteine as the starting reactant (de Clercq, 1997). We assumed a yield of 50% due to the multiple reaction steps with potential for loss and differences in molecular weight. The yield could be potentially lower due to inefficiencies in production. Adequate data were not available to estimate inputs/outputs during the multiple step conversion process. Cysteine can be produced microbially utilizing glucose as a feedstock and the embedded resources were determined utilizing the method described in the microbial methods section and appendix B (Martin-Dominguez et al., 2018).

The embedded resources associated with mass of glucose was then attributed to biotin. The reported embedded resource estimates for biotin production should be considered a minimum due to unavailability of input/output data related to energy and material flows during the production.

#### Sodium selenite

The following production route was chosen for sodium selenite, however sodium selenite can be prepared utilizing other methods (Brauer, 1963). This production route was chosen to minimize the additional compounds required for sodium selenite production. After stoichiometric calculations were complete, ecoinvent datasets associated with selenium production (selenium, soda ash, sulfur dioxide and hydrogen chloride) were utilized to quantify the some of the embedded resources in sodium selenite production. Sodium hydroxide, soda ash, sulfur dioxide and hydrogen chloride are utilized in production of this component and multiple E8/B9 components (See appendix G for complete list). Ecoinvent datasets were not available for sodium selenite and selenium dioxide production.

Equation A2. Selenium dioxide production

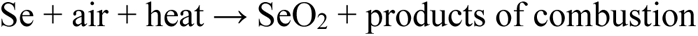

Assumption: No Se is lost during conversion of Se to SeO_2_. Theoretical yield is utilized. Energy usage is not calculated for stoichiometric equations.

Equation A3. Sodium selenite production

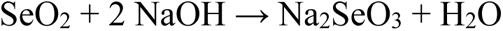

Assumption: Theoretical yield is utilized. Energy usage is not calculated for stoichiometric equations.

#### Cuperic Sulfate

Ecoinvent datasets were utilized to estimate material and energy flows associated with cuperic sulfate production (copper sulfate, copper oxide and copper cathode). Sulfuric acid is utilized in production of this component and multiple E8/B9 components (See appendix G) for related ecoinvent datasets. Copper cathode production data set includes embedded resources from copper mining.

#### Ferric Nitrate

Ferric nitrate can be prepared by adding nitric acid to iron pellets, powder or scrap iron (National Center for Biotechnology, 2021). After stoichiometric calculations were complete, ecoinvent datasets (Iron mining and beneficiation, iron pellet and nitric acid) were utilized to quantify the some of the embedded resources and outputs in ferric nitrate production. Ecoinvent data set was not available for ferric nitrate production.

Equation A4. Ferric nitrate production

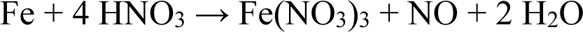

Assumption: Theoretical yield is utilized.

#### Ferrous sulfate

Ferrous sulfate is generally produced as a by-product of steel manufacture (Wildermuth et al., 2000). The utilized ecoinvent dataset (iron sulfate) accounts for this and only the embedded resources from additional refining is accounted for in the dataset and our model.

#### Folic acid

Chemical synthesis of folic acid largely occurs in an inexpensive, one-pot process (Mair et al., 2019). The economic viability of the chemical synthesis of folic acid largely prevents other production methods such as microbial fermentation or refinement from raw materials. Adequate data was not available to estimate embedded energy, but wastewater production was estimated in be 250-300 kg of wastewater for each kg of folic acid produced(Bryne, 2015).

#### Zinc Sulfate

Ecoinvent data sets were utilized to estimate embedded resources and outputs in zinc sulfate production (zinc sulfate and zinc oxide). Sulfuric acid is utilized in production of this component and multiple E8/B9 components (See appendix G). The embedded resources for zinc scrap are not included due to being a by-product of other production process and iron scrap being utilized as a stand in the dataset.

#### Phenol red

Phenol red (phenolsulfophthalein) can be obtain by the condensation of o-sulfobenzoic acid anhydride with phenol (Gessner & Mayer, 2011). Sulfobenzoic acid (mostly m-sulfobenzoic acid) derived from benzoic acid and sulfuric acid is used as a substitute for pure o-sulfobenzoic acid (Reese, 1932). Stochiometric calculations were conducted to determine the mass of compound needed when information was unavailable in ecoinvent datasets. Ecoinvent datasets were utilized to estimate embedded resources and outputs in phenol red production (phenol, benzoic acid, toluene, and benzene). Sulfuric acid, propylene, oxygen, in ground natural gas and crude oil are utilized in production of this component and multiple E8/B9 components (See appendix G). Ecoinvent datasets were unavailable for phenol red and sulfobenzoic acid.

#### Putrescine-2HCl

Putrescine and HCl required mass and embedded resources were determined utilizing stochiometric calculations and ecoinvent datasets. Putrescine can be produced utilizing acrylonitrile and hydrogen cyanide in the presence of tertiary amine and with subsequent hydrogenation (Broadwith, 2011). Ecoinvent datasets were utilized to estimate the material flows in Putrescine-2HCl (Acrylonitrile, HCl, hydrogen cyanide). Both acrylonitrile and HCl are utilized in production of this component and multiple E8/B9 components (See appendix G). Ecoinvent datasets were unavailable for putrescine and putrescine-2HCl.

#### Glycine

Ecoinvent data sets were utilized to estimate energy and material flows during glycine production (glycine, chloroacetic acid, acetic acid, and carbon monoxide). Ammonia, chlorine (liquid), sodium hydroxide and methanol are utilized in the production of this component as well as multiple E8/B9 components (See section appendix G).

#### Choline Chloride

Choline chloride is produced via reaction between hydrochloric acid, trimethylamine and ethylene oxide (equation A5) (Johnson Matthey Davy Technologies, 2014). Ecoinvent datasets were utilized to estimate embedded resources (ethylene oxide and trimethylamine). Hydrochloric acid, ethylene oxygen, ammonia, methanol, in ground natural gas, and crude oil are utilized in the production of this component as well as multiple E8/B9 components (See appendix G). An ecoinvent dataset was unavailable for choline chloride.

Equation A5. Choline chloride production

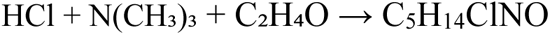

Assumption: Theoretical yield is utilized.

#### Niacinamide

Niacinamide can be produced utilizing microbial synthesis with precursor compounds being produced chemically (Shimizu et al., 2000; Z. Wang et al., 2017). Niacinamide can be produced via microbial conversion of 3-cyanopyridine to niacinamide with a 94.5% yield (Z. Wang et al., 2017). 3-cyanopyridine can be produced via the ammoxidation of 3-methylpryidine (equation A6) (Shimizu et al., 2000). Ecoinvent datasets were utilized to estimate embedded resources for niacinamide (3-methylprydine and acrolein). Ammonia, oxygen and propylene are utilized for the production of this component as well as multiple E8/B9 components (See appendix G).

Embedded resources for niacinamide and 3-cyanopyridine production are not accounted.

Equation A6. 3-cyanopydrine production

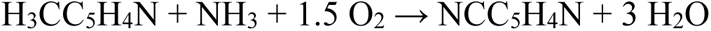

Assumptions: Theoretical yield is utilized. Stoichiometric calculation does not account for high density cell growth in bioreactor.

#### Thiamine hydrochloride

Thiamine hydrochloride can be manufactured by combining thiamine chloride with one molar equivalent of hydrochloric acid (ChEBI, 2021). Thiamine production process was then utilized as substitute for thiamine chloride production process and acrylonitrile was utilized as the starting material for thiamine production (Létinois et al., 2020). Acrylonitrile and hydrochloric acid production was then utilized to estimate embedded resources in thiamine hydrochloride production. The embedded resources utilized during thiamine and the intermediates production processes are not included in the assessment.

### Appendix E- Solvay

#### Sodium bicarbonate

Sodium bicarbonate production can be integrated into a soda ash production utilizing the Solvay process (European Commission, 2007b). To produce one ton of sodium bicarbonate approximately 0.7 tonnes of raw soda ash and 550 kg of CO_2_ (approximately 53% of the CO2 is released to atmosphere) are ulitilzed (European Commission, 2007b). Equation A7 illustrates the theoretical mass balance to produce sodium bicarbonate. The CO_2_ utilized for soda ash is considered to be a product of combustion and embedded resources for the CO_2_ is not accounted.

Equation A7. Theoretical sodium bicarbonate production

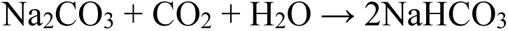

#### Calcium chloride

Calcium chloride is assumed to be produced from the Solvay process and soda ash. The calcium chloride is assumed to dried and the embedded resources for liquor production and drying processes are accounted for (European Commission, 2007a). The drying process can produce a product that is 75-82% flake or 100% prills (European Commission, 2007a). The 100% prills production was assumed to be the product utilized for E8/B9 production. Table A3 provides estimates for embedded energy and water in use for calcium chloride production.

**Table A3.** Energy and water inputs for calcium chloride prills (100% w/w) production from soda process

|  |  |
| --- | --- |
| Thermal energy for liquor production (GJ/ton) | 7-9 |
| Electrical energy for liquor production (GJ/ton) | 0.3-0.5 |
| Cooling water for liquor production (m<sup>3</sup>/ton) | 40 |
| Thermal energy for prills production (GJ/ton) | 7-9 |
| Electrical energy for prills production (GJ/ton) | 0.6-0.8 |
| Cooling water for prills production (m<sup>3</sup>/ton) | n/a |

\*Table developed from European Union reference document (European Commission, 2007a)

#### Sodium Phosphates

Sodium phosphates (monobasic and dibasic) are estimated to require approximately the same embedded resources and are not delineated between in embedded resource summation. Sodium phosphate production requires the use of soda ash or sodium hydroxide for production and are categorized as a product of the Solvay process due to soda ash being a major reactant. Ecoinvent datasets (sodium phosphate, purified phosphoric acid, phosphoric acid (fertilizer), quicklime, and beneficiated phosphate rock) were utilized to quantify the embedded resources in the sodium phosphate production process. Soda ash, sulfuric acid and crushed limestone are utilized for this component as well as multiple E8/B9 components (See section xx for related ecoinvent datasets).

### Appendix F- Potash

#### Potassium Chloride

Potassium chloride is assumed to be produced from a mining operation and the ecoinvent dataset starts at the extraction at mine and ends with production of 1 kg of potassium chloride.

#### Magnesium Chloride

Magnesium chloride can be produced from a variety of sources including salt lakes, underground brines, residual brines from the potash industry (European Commission, 2007a). The embedded resources were estimated utilizing industry supplied data as an estimation for embedded energy and water (Table A4) (Compass minerals, 2016). Energy and water usage calculated using reported water and energy intensities of products (Compass minerals, 2016).

**Table A4.** Embedded water and energy in magnesium chloride production

|  |  |
| --- | --- |
| Energy (MJ/kg) | $4.9 \times 10^{-1}$ |
| Water (m<sup>3</sup>/kg) | $4.7 \times 10^{-3}$ |

#### Magnesium Sulfate

Magnesium sulfate is assumed to be produced from a mining operation and the reported embedded resources and outputs in the ecoinvent dataset starts at the extraction at mine and ends with production of 1 kg of magnesium sulfate.

### Appendix G- List of components used in production of multiple E8/B9 components

**Table A5.**
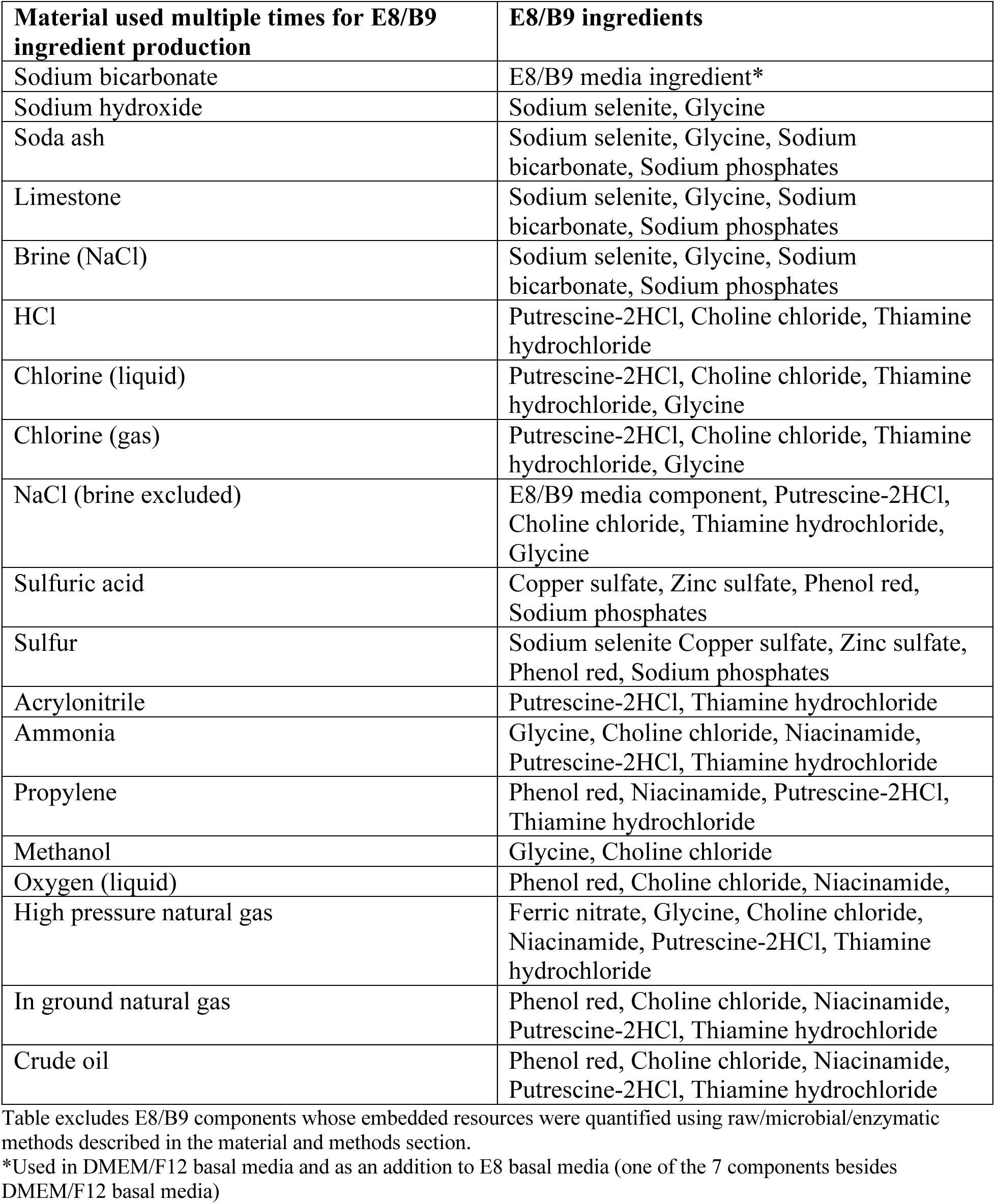
Materials used to produce more than one E8/B9 ingredient

| **Material used multiple times for E8/B9 ingredient production** | **E8/B9 ingredients** |
| --- | --- |
| Sodium bicarbonate | E8/B9 media ingredient* |
| Sodium hydroxide | Sodium selenite, Glycine |
| Soda ash | Sodium selenite, Glycine, Sodium bicarbonate, Sodium phosphates |
| Limestone | Sodium selenite, Glycine, Sodium bicarbonate, Sodium phosphates |
| Brine (NaCl) | Sodium selenite, Glycine, Sodium bicarbonate, Sodium phosphates |
| HCl | Putrescine-2HCl, Choline chloride, Thiamine hydrochloride |
| Chlorine (liquid) | Putrescine-2HCl, Choline chloride, Thiamine hydrochloride, Glycine |
| Chlorine (gas) | Putrescine-2HCl, Choline chloride, Thiamine hydrochloride, Glycine |
| NaCl (brine excluded) | E8/B9 media component, Putrescine-2HCl, Choline chloride, Thiamine hydrochloride, Glycine |
| Sulfuric acid | Copper sulfate, Zinc sulfate, Phenol red, Sodium phosphates |
| Sulfur | Sodium selenite Copper sulfate, Zinc sulfate, Phenol red, Sodium phosphates |
| Acrylonitrile | Putrescine-2HCl, Thiamine hydrochloride |
| Ammonia | Glycine, Choline chloride, Niacinamide, Putrescine-2HCl, Thiamine hydrochloride |
| Propylene | Phenol red, Niacinamide, Putrescine-2HCl, Thiamine hydrochloride |
| Methanol | Glycine, Choline chloride |
| Oxygen (liquid) | Phenol red, Choline chloride, Niacinamide, |
| High pressure natural gas | Ferric nitrate, Glycine, Choline chloride, Niacinamide, Putrescine-2HCl, Thiamine hydrochloride |
| In ground natural gas | Phenol red, Choline chloride, Niacinamide, Putrescine-2HCl, Thiamine hydrochloride |
| Crude oil | Phenol red, Choline chloride, Niacinamide, Putrescine-2HCl, Thiamine hydrochloride |

Table excludes E8/B9 components whose embedded resources were quantified using raw/microbial/enzymatic methods described in the material and methods section.
\*Used in DMEM/F12 basal media and as an addition to E8 basal media (one of the 7 components besides DMEM/F12 basal media)

### Appendix H- Transportation

Ecoinvent datasets were utilized to account for the resources used for transportation across each components supply chain. These data sets were then entered into OpenLCA and the LCIA methods software package was utilized to determine the environmental impacts of transportation across the supply chain. Additional information related to the transportation of E8/B9 can be found in the following appendix subsections.

#### Raw ingredients

##### Glucose

Each market was utilized for each stage of production (dry maize to glucose). The total embedded energy related to corn/glucose transport was determined then the embedded energy used per kg of glucose was determined.

##### Linoleic acid

Refined vegetable oil market as a substitute in the ecoinvent dataset. The corn transport was not accounted for on based on the assumption that wet milling will be utilized and is onsite.

#### Microbial-Yield and titer

Glycine which is an amino acid produce in bulk and the market is reported in ecoinvent will be utilized as a substitute for the transportation. The total required glucose will also be accounted for utilizing transportation per kilogram of glucose.

#### Enzymatic, Aspartic acid

Glucose and it’s embedded resources utilized for transportation was utilized to estimate the embedded energy related to aspartic acid transportation.

#### Chemical, Biotin

The embedded transportation energy from cysteine and glucose transport was utilized to estimate embedded transportation energy.

### Appendix I- Sensitivity analysis

DMEM/F12 basal medium was found to be the most environmentally impactful component of both E8 and B9af in all impact categories. To further understand the drivers of these impacts, an additional sensitivity analysis was conducted to determine which of the DMEM/F12 basal medium components (>50) most influenced its environmental impact. The sensitivity analysis indicates that the glucose input is the most environmentally impactful DMEM/F12 component. This is due to the glucose concentration being orders of magnitude greater than most other inputs except sodium chloride and HEPES. A 25% increase in glucose concentration changed each TRACI 2.1 output by 6-20% and each cumulative energy demand output by 6-12% (See appendix figures A1.0 and A2.0). A 25% increase sodium bicarbonate or sodium chloride concentration increased some TRACI outputs by ∼3% and cumulative energy demand outputs by ∼2%. Looking at broader categories of inputs, a 25% increase in all amino acids increases CED and TRACI 2.1 outputs by ∼15%. It should also be noted that HEPES concentration is greater than glucose concentration, but the HEPES environmental impacts are not accounted for due to a lack of manufacturing data.

**Figure A1.**
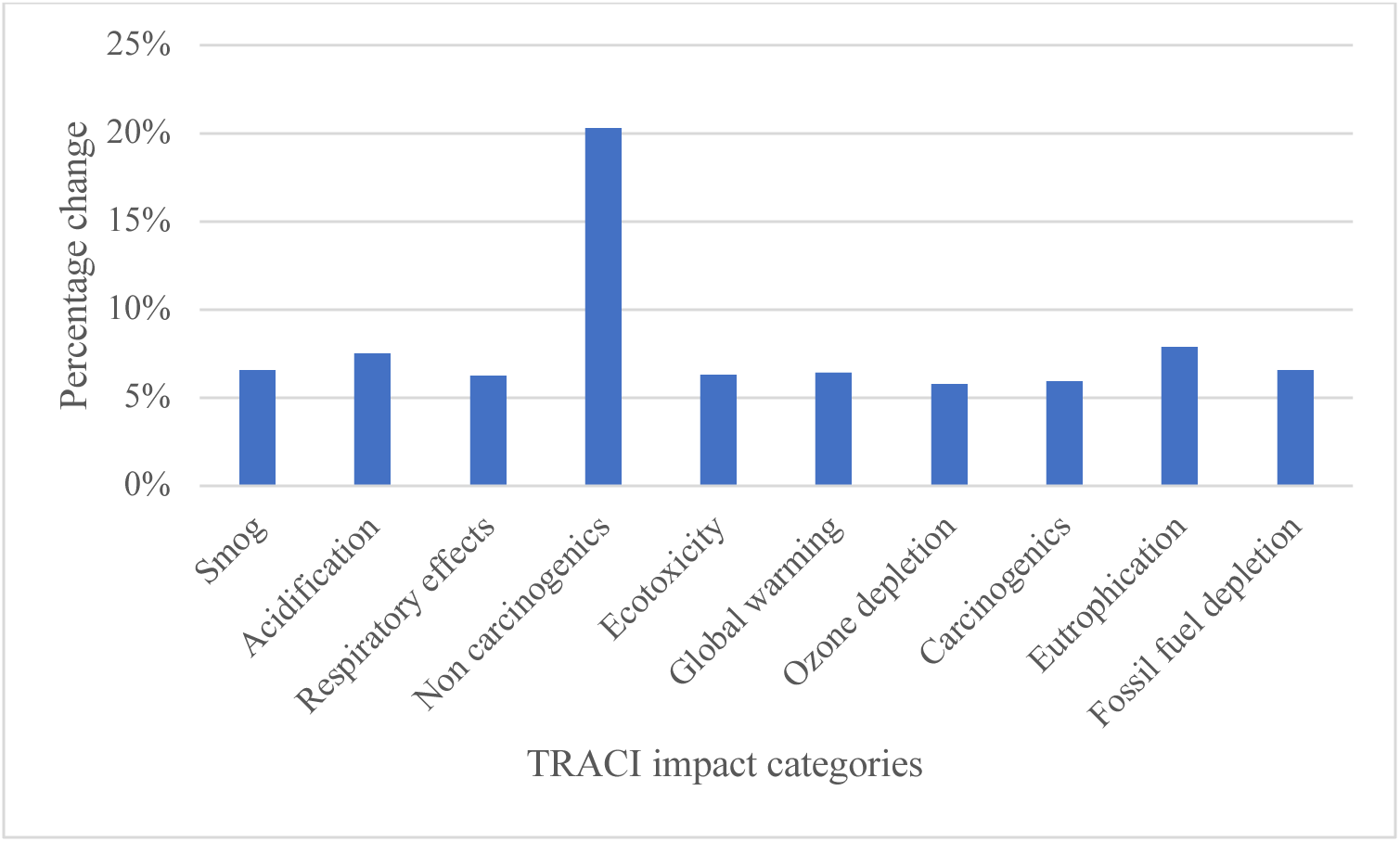
Percentage of change in each TRACI impact category from a 25% increase in DMEM/F12 glucose concentration

**Figure A2.**
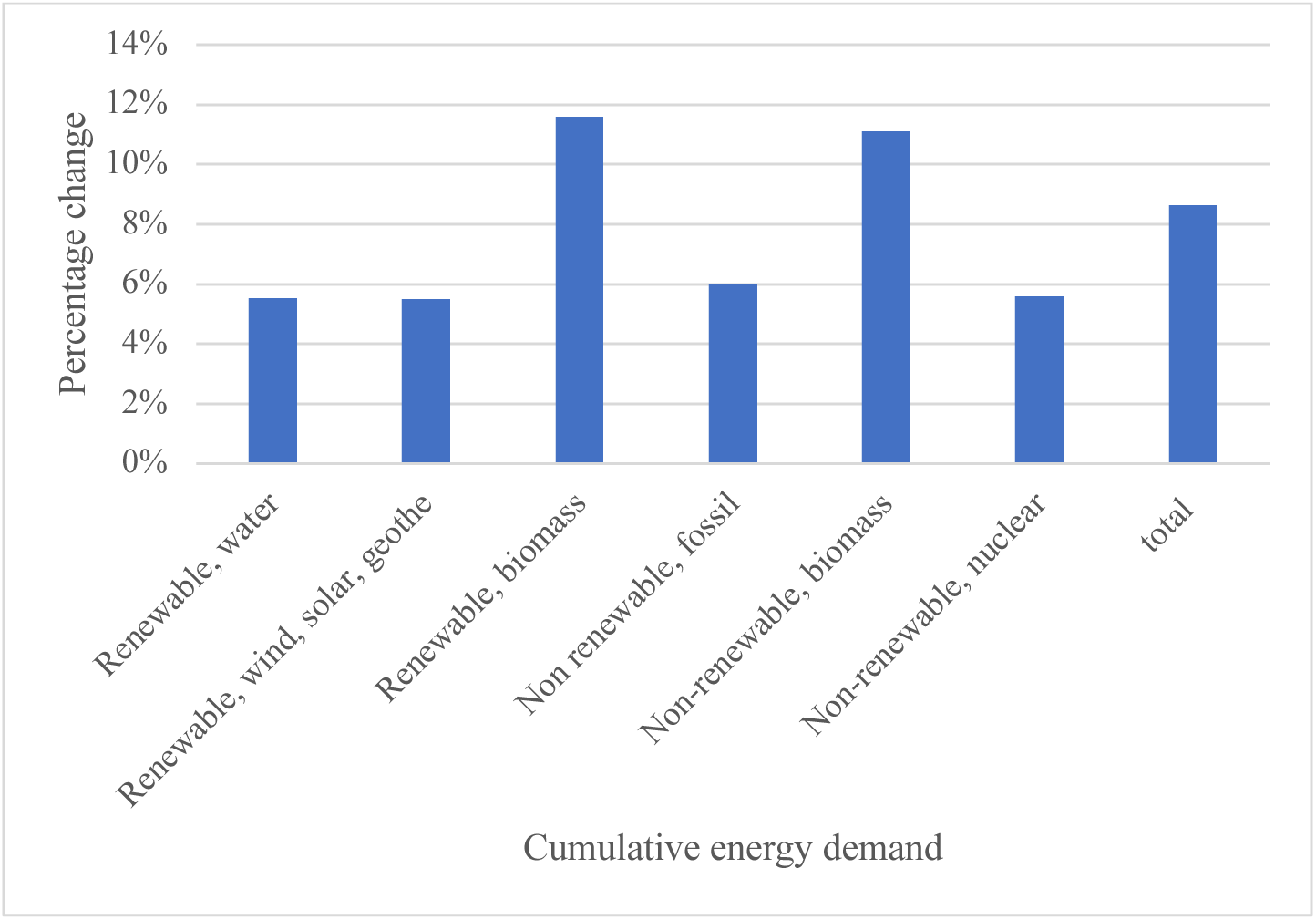
Percentage of change in each cumulative energy demand category from a 25% increase in DMEM/F12 glucose concentration

